# Fractal: Towards FAIR bioimage analysis at scale with OME-Zarr-native workflows

**DOI:** 10.64898/2026.03.05.709921

**Authors:** Joel Lüthi, Lorenzo Cerrone, Tommaso Comparin, Max Hess, Ruth Hornbachner, Adrian Tschan, Gustavo Quintas Glasner de Medeiros, Nicole A. Repina, Loredana K. Cantoni, Fabio D. Steffen, Jean-Pierre Bourquin, Prisca Liberali, Lucas Pelkmans, Virginie Uhlmann

## Abstract

The rapid growth in microscopy data volume, dimensionality, and diversity urgently calls for scalable and reproducible analysis frameworks. While efforts on the open OME-Zarr format have helped standardize the storage of large microscopy datasets, solutions for standardized processing are still lacking. Here, we introduce two complementary contributions to address this gap: 1) the Fractal task specification, defining OME-Zarr processing units that can interoperate across computational environments and workflow engines, and 2) the Fractal platform, using this specification to enable scalable and modular OME-Zarr-native analysis workflows. We demonstrate their use across diverse biological research data, including terabyte-scale multiplexed, volumetric, and time-lapse imaging. In a clinical setting, we show that Fractal workflows achieve near-identical quantification of millions of cells across independent deployments, demonstrating the reproducibility required for translational applications. With its growing community of contributors, the Fractal ecosystem provides a foundation for FAIR microscopy image analysis relying on open file formats.

## Introduction

Recent breakthroughs in artificial intelligence (AI) have made the systematic use of bioimages as quantitative data a reality. AI technologies initially developed for computer vision applications and subsequently adapted for microscopy offer unprecedented opportunities to automate the analysis of large, multidimensional datasets^1,2^ and even allow prediction of patterns that have not been seen during training^3,4^. The past decade has seen considerable effort to democratize these AI methods through user-friendly interfaces, exemplified by ImageJ/Fiji^5^, napari^6^, and ilastik^7^, among many others. However, ever-growing dataset sizes introduce substantial computational and engineering challenges, with problems that are manageable at megabyte scales becoming insurmountable at terabyte scales. Compounding this challenge, microscopy has to deal with an extreme data diversity: acquisition systems produce images with different formats, dimensions, and sizes, with new modalities continuously emerging. While AI-ready microscopy can unlock new research frontiers, achieving this readiness presents a formidable engineering challenge: the immense diversity of formats, the ever-increasing variety of analysis algorithms, and the sheer scale of data, where semantic information spans multiple dimensions, requires the development of novel solutions^8^.

Clearly, specialized approaches reinvented for each modality cannot solve the global big data challenge that microscopy imaging is facing. Instead, data diversity must be embraced through standardized frameworks that abstract modality-specific differences and enable the development of reusable, general analysis methods tailored to the needs of bioimaging^9^. Towards that end, international community efforts have progressed to define standards that support Findable, Accessible, Interoperable, and Reusable (FAIR^10^) microscopy. Over two decades, metadata standards for bioimaging have successfully been established through the Open Microscopy Environment (OME) data model^11^. Beyond metadata, international community efforts have initiated crucial discussions on next-generation file formats (NGFF^12^) that standardize both the metadata (i.e., image content and acquisition parameters) and the data themselves (i.e., actual images) in scalable ways. These efforts culminated in the OME-Zarr implementation proposal^13^, combining Zarr, a general-purpose format for large n-dimensional arrays, with community-defined bioimage metadata specifications. The 2024 OME-NGFF Challenge^14^ further demonstrated the potential of this format as a unifying standard, with the international community making hundreds of terabytes of diverse microscopy imaging data globally accessible within a few months. Ever since, the OME-Zarr community has continued to expand rapidly, with worldwide contributions to the specification as well as to the development of image viewers^15–17^ and tooling^18,19^. With image format standardization across modalities and scales now within reach, the time is ripe to leverage OME-Zarr for large-scale, FAIR microscopy analysis. Yet, standardized data alone is insufficient: making microscopy image analysis interoperable requires defining how processing operations should read, transform, and write OME-Zarr files across different computational environments, as well as how workflows can be run in a scalable and reproducible fashion.

In this paper, we introduce two complementary contributions: first, the Fractal task specification, which establishes a strategy for FAIR processing on OME-Zarr, and second, the Fractal platform, which demonstrates how this specification enables the design and execution of OME-Zarr-native image analysis workflows in practice. The Fractal task specification is built on the idea that image analysis operations (referred to as “tasks”) operating from on-disk OME-Zarr to on-disk OME-Zarr constitute a powerful interoperability unit. This disk-to-disk schema enables task execution across fundamentally different workflow engines and in distinct computational environments. The Fractal platform then offers as a concrete demonstration of the utility of this specification and provides a workflow management framework with a user-friendly, modular approach to large-scale bioimage analysis. Growing interest in microscopy analysis workflows has spurred extensive efforts to democratize workflow orchestration platforms such as Nextflow^20^, Snakemake^21^, Galaxy^22^, and others for microscopy applications. While these platforms offer powerful general-purpose orchestration capabilities, their support for OME-Zarr however remains limited, creating an opportunity for an interoperable specification that would bridge these ecosystems. Together, our two contributions represent, to the best of our knowledge, the first proposal for an interoperable microscopy image processing strategy, built entirely around the OME-Zarr format. As such, they offer a path toward OME-Zarr adoption across the broader workflow orchestration community by enabling the development of processing units that integrate across existing and future frameworks.

Our contributions expand the definition of FAIR microscopy imaging from data management to analysis: the Fractal task specification and platform make it possible to develop custom processing modules for specific applications that are standardized through their reliance on OME-Zarr. Over 100 publicly-available analysis tasks implemented following the Fractal task specification are currently available, with this collection continuously expanding through community contributions. These include a broad set of (i) conversion tasks to transform vendor-specific proprietary image formats into OME-Zarr, (ii) tasks for image-based registration across multiplexing experiments and for image stitching, (iii) tasks for object segmentation leveraging both classical methods as well as various popular deep learning models, and (iv) tasks for measurements extracting quantitative descriptors of segmented objects, including morphology and image intensity among others. We demonstrate the applicability of the Fractal task specification and platform for the analysis of diverse challenging biological imaging data, including terabyte-scale multiplexed, volumetric, and time-lapse imaging, as well as in a clinical setting where reproducibility and traceability are paramount. Looking forward, this kind of standardized, interoperable approach to microscopy image processing has the potential to transform how the life sciences community collaborates, as it facilitates the rapid global deployment and adoption of breakthrough analysis methods.

## Results

### The Fractal task specification: OME-Zarr-native interoperable processing units

The OME-Zarr specification offers a solid basis for the structured storage of both raw and derived microscopy images^13^. Built on the Zarr format, a cloud-compatible storage container for N-dimensional arrays that is used across disciplines ranging from physics to astronomy and Earth science^23^, OME-Zarr allows storing larger-than-memory image data of up to 5 dimensions in chunked Zarr stores both at their full resolution as well as at different downsampled resolutions that allow for easier visualisation (**Figure 1A**). By recording well-defined metadata on the nature of each resolution and channel, OME-Zarr makes it easier to interpret large image data. Additionally, derived data such as segmentation masks can be stored in the same OME-Zarr container as the original images, thereby linking raw and derived data. Here, we propose to enrich the OME-Zarr containers with the ability to save tabular data in diverse stores such as AnnData^24^, Apache Parquet, CSV, or JSON, directly in the OME-Zarr container. This offers an extensible framework for storing feature measurements, coordinates of region of interest (ROI), tabular metadata, and other tabular formats together in a single OME-Zarr file.

**Figure 1.**
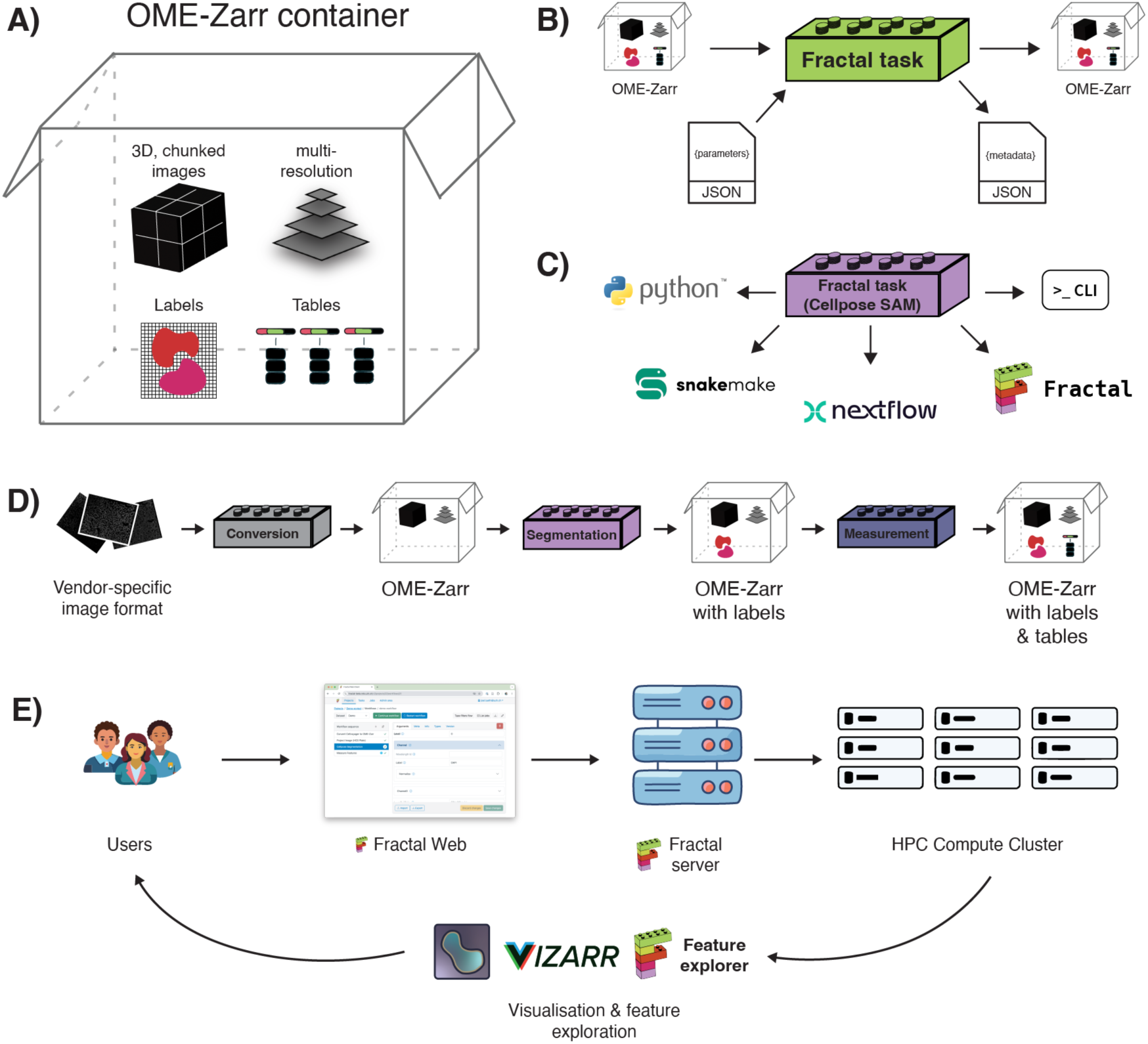
The Fractal task specification and the Fractal platform enable the interoperable processing of image data in the open OME-Zarr file format. **A** In addition to the image data themselves (both at full resolution and in downsampled pyramid levels), metadata, and derived label data, our OME-Zarr containers can also contain tabular data for different table subtypes, making them self-contained units for image processing. **B** Image processing modules implemented following our proposed Fractal task specification take OME-Zarrs and parameters in JSON format as input through the command line interface, and output data in the OME-Zarr format with accompanying JSON metadata. **C** The interoperability of the Fractal task specification is demonstrated with the example of a cellpose SAM task, which can be directly used through Python, command line interface, Snakemake, Nextflow, or the Fractal platform. **D** Data standardization through OME-Zarr allows chaining tasks across computational environments and combining tasks from multiple developers into customized analysis workflows. **E** Building upon our definition of standardized OME-Zarr containers and processing tasks, researchers can be empowered to build, execute, and monitor large-scale image analysis workflows processing terabytes of image data on HPC clusters through the web interface of the Fractal platform.

To enhance the interoperability of processing units developed to operate on standardized OME-Zarr containers, we introduce the Fractal task specification, which defines how computational units must be implemented to process OME-Zarr files and be reusable in various computational environments. The Fractal task specification defines processing units as command-line executables that load data from OME-Zarr containers, process them, and write results into the same or new OME-Zarr containers (**Figure 1B**). Parameters to configure processing tasks are provided as JSON files, and tasks can generate additional output metadata in the JSON format to specify any newly created OME-Zarr containers as well as their associated metadata. We illustrate how the Fractal task specification facilitates reusability through an example implementing the recent cellpose SAM model^25^. The process of creating a task performing instance segmentation on OME-Zarr files relying on the cellpose SAM model is facilitated through the use of templates, and the resulting task can straightforwardly be used in a Python script, a bash script via command line interface, a Nextflow workflow, or a Snakemake workflow (**Figure 1C**, additional details available at https://github.com/fractal-analytics-platform/fractal-task-interoperability). Importantly, these different ways of running the task do not require any modification of the task code itself. In addition to this example, the broad reusability of processing modules implemented to follow the Fractal task specification has been further explored by the community for other segmentation algorithms^26^, demonstrating the potential of this approach to enable interoperable microscopy image analysis.

As task inputs and outputs follow the standardized OME-Zarr containers format, tasks can be arbitrarily combined into workflows that apply a sequence of different processing steps to enrich OME-Zarr containers with new data (**Figure 1D**). Processing tasks following the Fractal task specification can be bundled into individual packages, each defining a collection of tasks that can be run in the same computational environment. A JSON manifest file defines the list of available tasks in a given package as well as their parameters and additional metadata, such as docstrings and default computational requirements. At the time of writing, we and other members of the bioimage analysis community have made 16 task packages containing over 100 individual tasks publicly available under permissive, open-source licenses (full list available at https://fractal-analytics-platform.github.io/fractal_tasks). This community-driven approach increases the accessibility and findability of processing modules in the context of FAIR bioimage analysis workflows.

### The Fractal platform: OME-Zarr-native modular analysis workflows

The Fractal platform offers an OME-Zarr native approach to designing and executing workflows based on the Fractal task specification. It is designed as a federated approach to host an image analysis service accessible to scientists without programming expertise (**Figure 1E**). Once configured on a high-performance computing (HPC) infrastructure, the Fractal platform provides a web interface accessible through all common internet browsers to allow researchers to build workflows, manage their execution, and monitor their progress. As such, the Fractal platform provides an intuitive interface to interact with complex, scalable sequences of processing operations on OME-Zarr containers that can involve terabytes of image data on HPC clusters.

Fractal relies on several modular components. First, a backend server manages users, their access, and their workflows. Through a runner interface, the backend server talks to an HPC cluster to schedule workflows and monitor their progress. Finally, a web interface makes the service accessible in a no-code manner (**Supplementary Figure 1A**, **Supplementary Video 1**). As accessing data stores on HPC clusters is often not straightforward, the Fractal platform enables streaming access to the OME-Zarr containers it creates through a dedicated data service. This allows the interactive visualisation of image data and of their corresponding processing results through web-based viewers such as ViZarr^15^ (**Supplementary Figure 1B**) or locally-installed viewers such as napari^6^. Relying on napari plug-ins, selected regions of interests in images, labels, and tabular data can be visualized and further interactive data exploration can be carried out (**Supplementary Figure 1C**, **Supplementary Video 2**) on OME-Zarr containers that are available either locally (e.g., stored directly on a local hard drive or accessible on a mounted share) or remotely. The exploration of quantitative processing results saved as tables of measurements is facilitated by our custom Fractal feature explorer web dashboard based on the Streamlit framework (https://streamlit.io/), which loads feature data from OME-Zarrs and provides interactive filtering and plotting tools (**Supplementary Figure 1D**, **Supplementary Video 3**, additional details available at https://github.com/fractal-analytics-platform/fractal-feature-explorer). As a result, the Fractal platform provides a first example of an integrated no-code solution for large-scale bioimage data analysis using the open OME-Zarr standard, leveraging our proposed interoperable definition of processing tasks.

The Fractal platform is developed in the open and under a permissive open-source license, allowing federated deployments and decentralized operation in different institutional environments. While individual Fractal deployments must be carried out in close collaboration with local HPC and scientific computing service teams, we provide a public server for demonstration purposes, accessible at https://fractal-demo.mls.uzh.ch. Although not allowing direct workflow execution, this demonstration server makes it possible to explore the Fractal interface and inspect some of the workflows described in this paper, as well as to experience the ViZarr integration and the Fractal feature explorer.

### Static multiplexed image analysis workflows

We illustrate an OME-Zarr-based analysis workflow on a newly-acquired 10 TB, 14 multiplexing cycle 10-day cardiac differentiation time course assay imaged across 23 wells of a 96-well plate (**Figure 2A**). Although the dataset contains 3D images, the approximately 1 million cells mostly form a 2-dimensional cell layer (20480×19440 pixels in XY for only 19-42 Z-slices in any given well) resulting in “2.5D” images. We considered 11 timepoints (day 0 before differentiation to 10 days of differentiation) at a 24h interval, fixed and imaged on a Yokogawa Cell Voyager 7000 across all 4i multiplexing rounds^27,28^, staining for key differentiation markers in all wells to allow a quantification of the differentiation heterogeneity across the time course (**Figure 2B**). Each multiplexing cycle originally produced approximately 160,000 TIFF files with vendor-specific metadata specifying their position in the full 96-well plate. The processed OME-Zarr containers will be publicly available in the Bioimage Archive (*data submission in progress, link to be added in the final version of the manuscript*).

**Figure 2.**
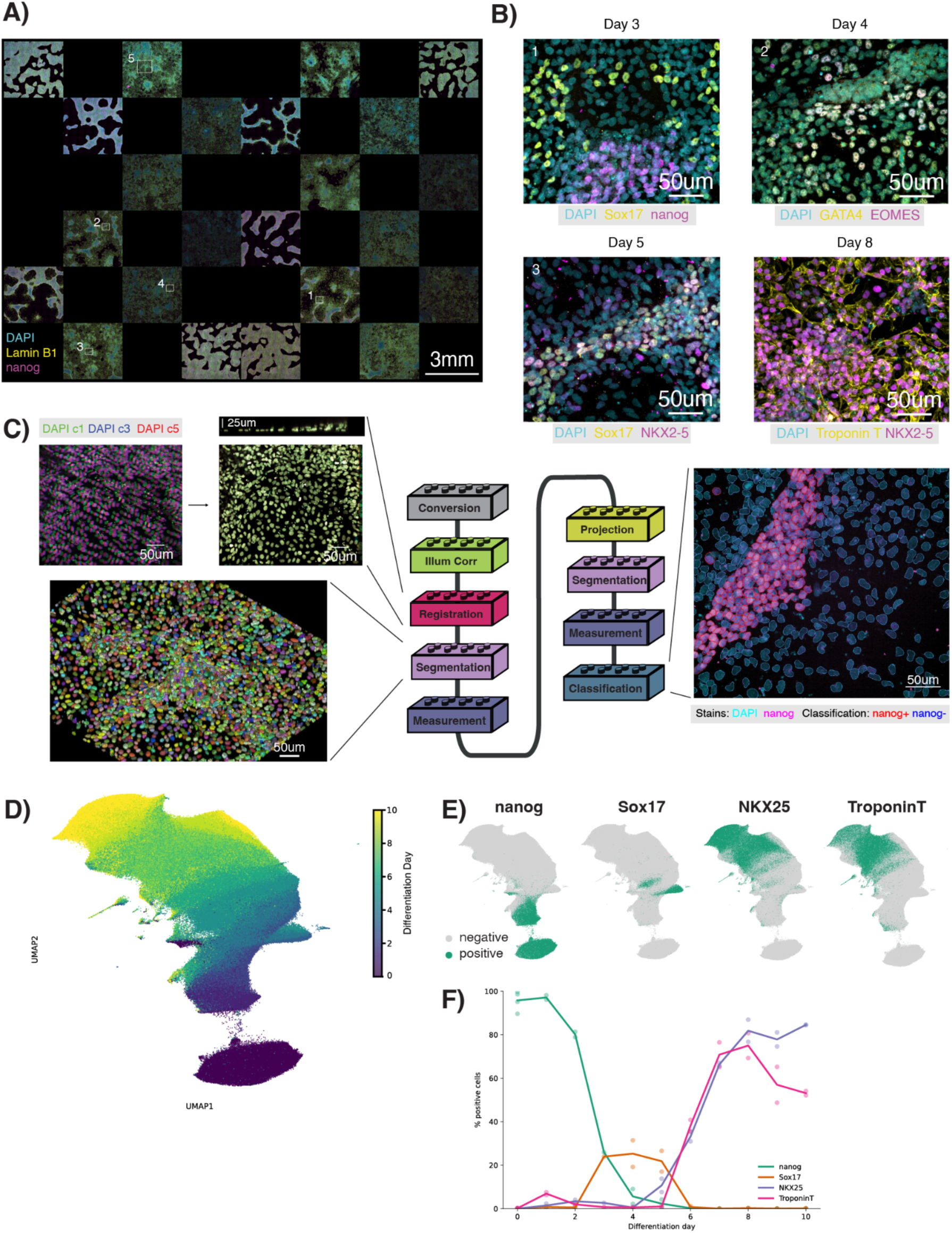
The Fractal task specification and the Fractal platform allow processing large multiplexed image datasets. **A-B** Using the Fractal platform, we converted a multiplexed cardiac differentiation time course of 10 TB of raw images acquired with a Yokogawa CV7000 Cellvoyager into OME-Zarr to obtain an overview of the whole differentiation course. **C** We designed a processing workflow including flatfield correction, 3D image registration, 2D and 3D nuclear segmentation, measurements extraction, and cell type classification. The top-left images illustrates the overlap of 3 multiplexing cycles before and after registration. The classification results in the right panel are shown as red (nanog+) vs. blue (nanog-) outlines. **D** With this analysis, we were able to relate stainings across multiplexing cycles and to generate highly multiplexed measurements of single cells across the cardiac differentiation trajectory. **E-F** The quantification carried out with our workflow recapitulates the gradual differentiation as well as the heterogeneity of differentiation outcomes, where only a part of the population of induced stem cells became early cardiomyocytes as measured by their expression of TroponinT starting at day 6 of differentiation.

We analyzed these data with a workflow starting with conversion to OME-Zarr, pre-processing to correct illumination biases and non-affine 3D registration between every multiplexing cycle and the reference cycle, individual 3D nuclei segmentation and quantification, maximum intensity projection, individual 2D nuclei segmentation and quantification, and cell type classification (**Figure 2C**). The resulting quantification of over 800,000 multiplexed cells after quality control with over 2500 measurements each, including morphology, intensity, and texture of the multiplexed markers across 10 days of cardiac differentiation, showcases the quantitative potential of large-scale image analysis (**Figure 2D**). Overlaying the classification results for 4 differentiation markers, we observed that the stemness marker nanog was only expressed in early differentiation days, the endodermal marker Sox17 was transitionally expressed in a clearly defined subgroup of cells between day 3 and 5 of differentiation, while cardiac differentiation markers such as NKX2-5 and Troponin T came up between day 5 and 6 respectively (**Figure 2E-F**). Our results recapitulate known outcomes of the cardiac differentiation protocol^29^, highlighting the capability of the Fractal framework to execute complex image analysis workflows across TB-scale multiplexed differentiation time courses.

### Static volumetric and multi-scale image analysis workflows

We further illustrate how the Fractal platform can be equally well used to process volumetric high-content and multiplexed imaging datasets of truly 3-dimensional biological structures. First, we re-processed a recently-published multiplexed protein imaging dataset (3D-4i) of zebrafish embryos^30^ using a newly-developed Fractal task collection (*abbott,* https://github.com/pelkmanslab/abbott) dedicated to 3D multiplexed imaging data (**Figure 3A**). The resulting workflow starts by conversion to OME-Zarr, followed by illumination correction, registration, various object segmentations, feature extraction, and finally classification (**Figure 3B**). As this data captures the spatiotemporal dynamics of the cell cycle and its relationship to the cellular state of the cells during zygotic genome activation, our analysis allows the study of self-organisation in developing multicellular systems across many zebrafish embryos (**Figure 3C**) while also providing quantitative volumetric information on their individual cells and nuclei (**Figure 3D**). The extracted measurements included multiplexed intensity and morphology readouts as well as spatial neighborhood information for each cell of the embryo. For a representative embryo of the 12th division cycle, we applied UMAP dimensionality reduction^31^ to generate a 2D projection of the high-dimensional, scale-crossing features extracted through the workflow (**Figure 3E**). Performing Leiden clustering^32^ on a graph constructed from the joint feature- and spatial-neighborhood space enables accurate identification of cell types present in the embryo, achieving approximately 90% of overlap with manual cell type annotations. The spatial organization of the resulting clusters can further be visualized by mapping the identified clusters back to the embryo in imaging space (**Figure 3F**). Notably, the observed separation of cells within the Deep Cell (DC) population likely reflects differences in cell cycle phases, as observed during early zebrafish development^30^. These findings illustrate the potential of Fractal to mine complex whole-mount multiplexed imaging data capturing information across multiple length scales and at high resolution, and to help unravel the drivers of self-organisation in developing multicellular systems. The processed OME-Zarr container will be publicly available in the Bioimage Archive (*data submission in progress, link to be added in the final version of the manuscript*).

**Figure 3.**
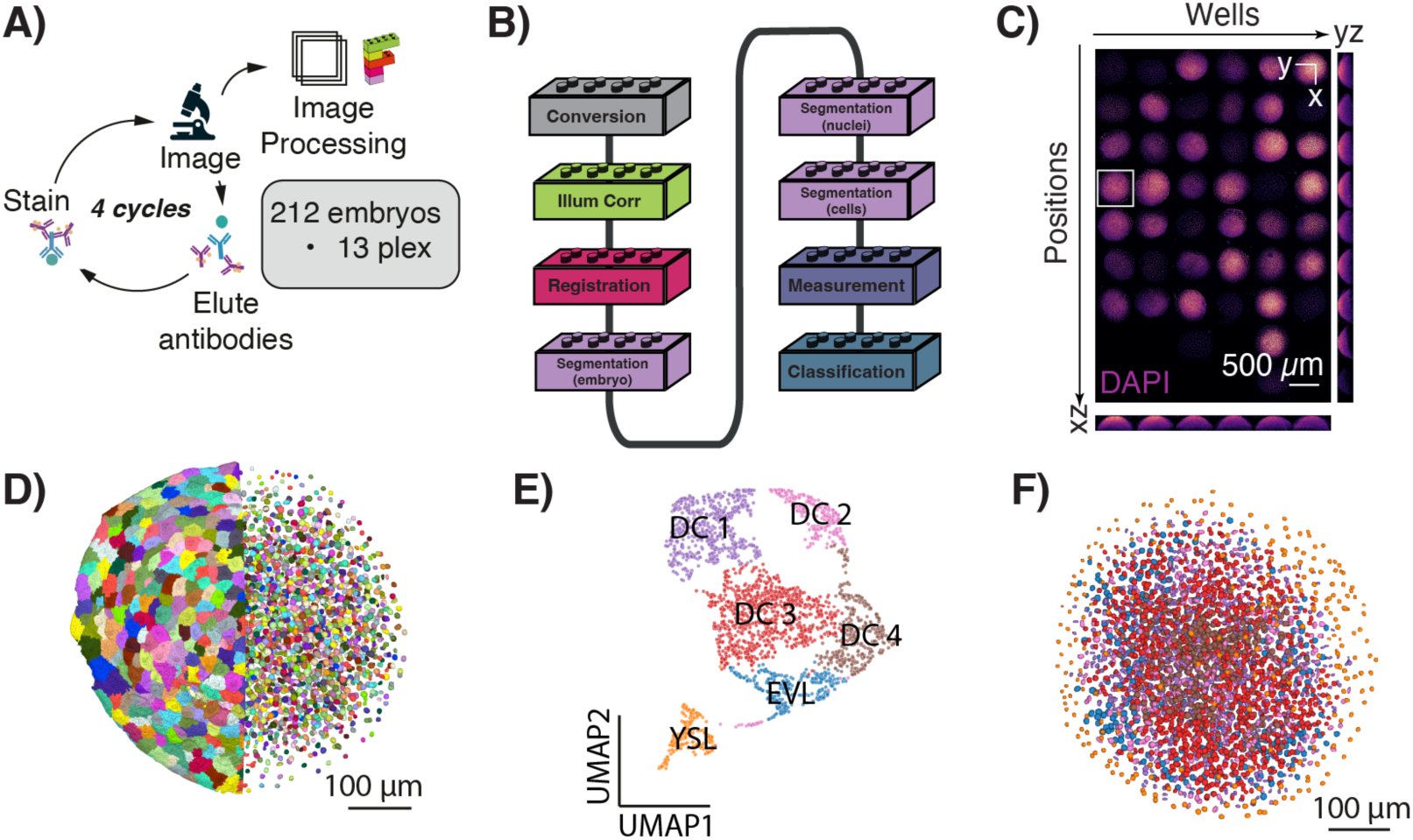
The Fractal task specification and the Fractal platform allow processing complex volumetric data. **A** Schematic overview of the 3D-4i workflow used to acquire a multiplexed time course of early zebrafish development. **B** Overview of the 3D multiplexed image processing workflow developed in the Fractal framework. **C** Maximum-intensity projections of a subset of zebrafish embryos acquired in the multiplexing experiment. Embryos are arranged from left to right by well position and from top to bottom by acquisition order. All embryos were stained with DAPI to label DNA. The white rectangle highlights the representative embryo shown in subsequent panels. **D** 3D rendering of segmentation results, depicting individual cells (left) and nuclei (right). **E** Spatially-aware Leiden clustering of the high-dimensional feature space obtained from the representative cycle 12 embryo, depicted as a 2D projection of the clusters using UMAP. Each cluster color corresponds to a distinct cell population, such as Deep Cell (DC) clusters 1-4, enveloping layer (EVL), and yolk syncytial layer (YSL). **F** Mapping of the identified clusters back onto the representative embryo in the original 3D imaging space.

Second, we used the Fractal platform to analyze molecular expression patterns, cell spatial location, and tissue-level shape properties in newly-acquired 3D multiplexed confocal imaging datasets of mouse small intestinal organoids (10-50 TB, **Figure 4A**). We developed a dedicated task collection to process these multiscale, multiplexed, and volumetric data (*scMultiplex*, https://github.com/fmi-basel/gliberal-scMultipleX) relying on a workflow starting with conversion to OME-Zarr, followed by 2D maximum intensity projection to identify parent objects (organoids) and link them across rounds. Each linked region was then processed in 3D to segment the surface epithelium of the organoid and convert it to a mesh format for further quantification and spatial analysis (**Figure 4B**). As a next step, further image segmentation was performed on the next level of organization (cells and nuclei) across the full volume of each organoid, and also converted to meshes (**Figure 4C**). This analysis resulted in a multiscale representation of thousands of organoids, from their tissue shape to their complete cellular composition (**Figure 4D**). Lastly, the 3D meshes were annotated by the molecular expression measured across multiplexed channels, which uncovers the relative spatial arrangement of cell types and signaling states in the system (**Figure 4E**). This workflow demonstrates how Fractal facilitates the processing of large image volumes on a per-region basis and on relevant multiplexing rounds for optimized memory usage and parallelization. It also illustrates how our framework enables the quantitative investigation of the role of molecular heterogeneity and cell spatial organization in collectively shaping tissue morphogenesis.

**Figure 4.**
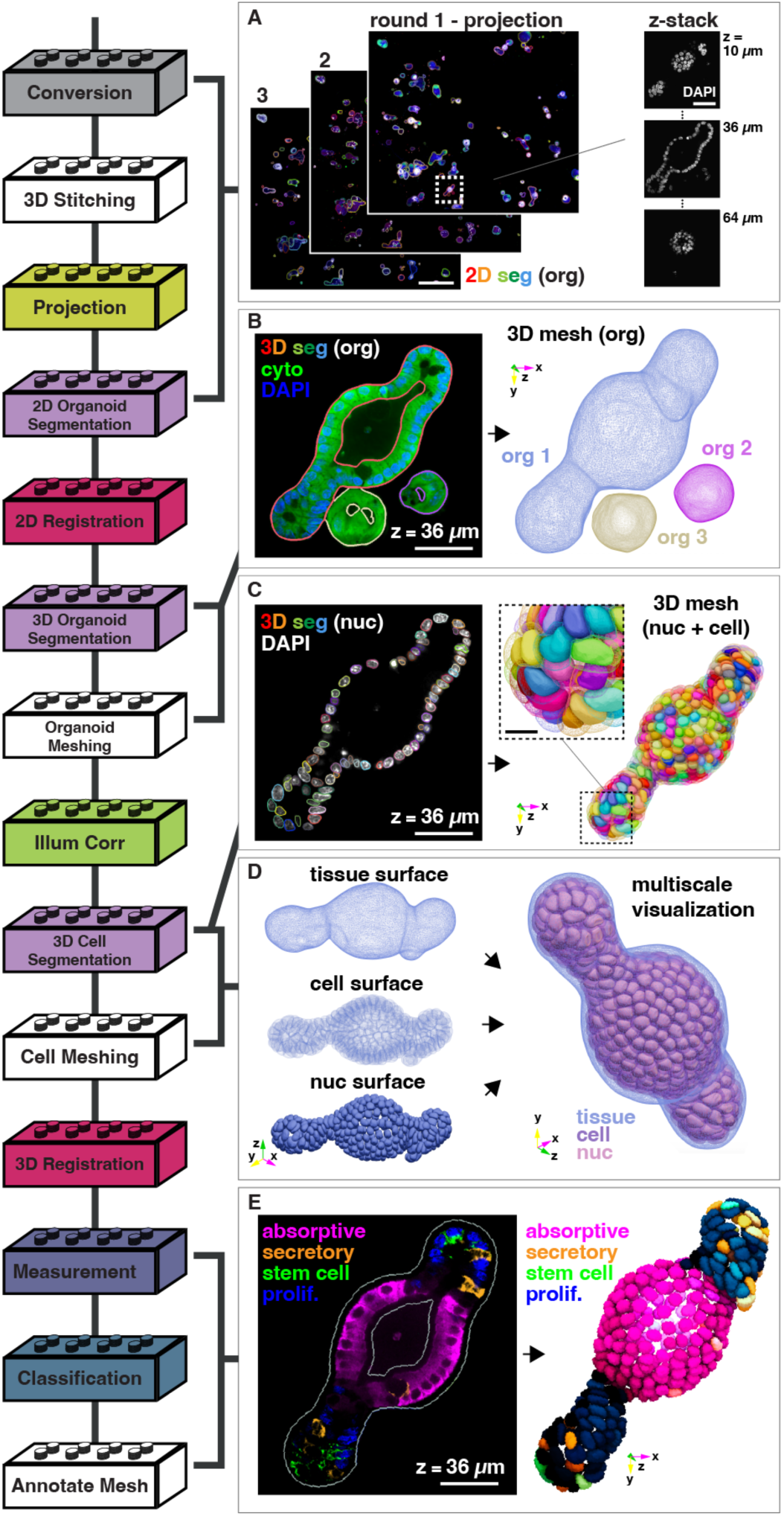
The Fractal task specification and the Fractal platform allow processing multiscale, multiplexed, and volumetric image data. **A** The raw spinning disk confocal microscope acquisition of multiple protein multiplexing rounds was converted to OME-Zarr and stitched across fields of view. The volumetric data were then projected to a 2D maximum intensity image for visualization and linking of organoids across multiplexing rounds (left, scale bar 500 µm). Each organoid was imaged with a fine Z-resolution that enabled 3D cell and tissue-level segmentation (right, scale bar 50 µm). **B** 3D tissue-level instance segmentations of cytoplasmic (cyto) fluorescence signal (left, scale bar 50 µm) were converted to organoid meshes of the surface epithelium (right). The color coding of the meshes corresponds to instance segmentation labels. **C** The 3D nuclear-level instance segmentation (left, scale bar 50 µm) was converted to a nuclear and cell mesh (right, scale bar 10 µm). The color coding of the meshes corresponds to instance segmentation labels. **D** Multiscale 3D visualization of the organoid tissue, cell, and nuclear composition. **E** Molecular expression of immunostained protein markers was used to identify cell types (left, scale bar 50 µm) and to annotate the nuclear mesh by molecular expression across registered multiplexing rounds (right). The color coding of the meshes is based on cell type classification, and color intensity corresponds to the 95% percentile intensity of the given protein marker

### Time-lapse volumetric image analysis workflows

In addition to static 3D data, Fractal can also be used to process live time-lapse volumetric images. To demonstrate this use-case, we created a minimal workflow for nuclear segmentation using the cellpose SAM task. We applied a simple workflow (**Figure 5A**) to a previously published dataset of mouse intestinal organoids developing from a single cell to a budding organoid over 110 hours of imaging^33^ (**Figure 5B**). The workflow segmented individual nuclei in all 667 3D timepoints (**Figure 5C**). Although the out-of-the-box performance of cellpose SAM was overall satisfactory, some nuclei appeared to be missed in degraded parts of the image in later timepoints (**Figure 5C**, white arrow) and apoptotic cells in the lumen resulted in false-positive detections (**Figure 5C**, red arrow). These limitations could be addressed in the future by extending the workflow with image restoration tasks and dedicated processing modules to segment the organoid and lumen, or by fine-tuning cellpose specifically to this kind of dataset. In spite of these segmentation errors, a quantitative analysis of the nuclei over time highlighted the growth of the organoid from a single to hundreds of cells (**Figure 5D**) and captured the synchronization of nuclear volume changes associated with the cell cycle in the first 50-60 hours of the time-lapse sequence (**Figure 5E**). These results exemplify how Fractal can facilitate the application of state-of-the-art deep learning models to complex, large, time-resolved datasets through its no-code interface. The processed OME-Zarr container will be publicly available in the Bioimage Archive (*data submission in progress, link to be added in the final version of the manuscript*).

**Figure 5.**
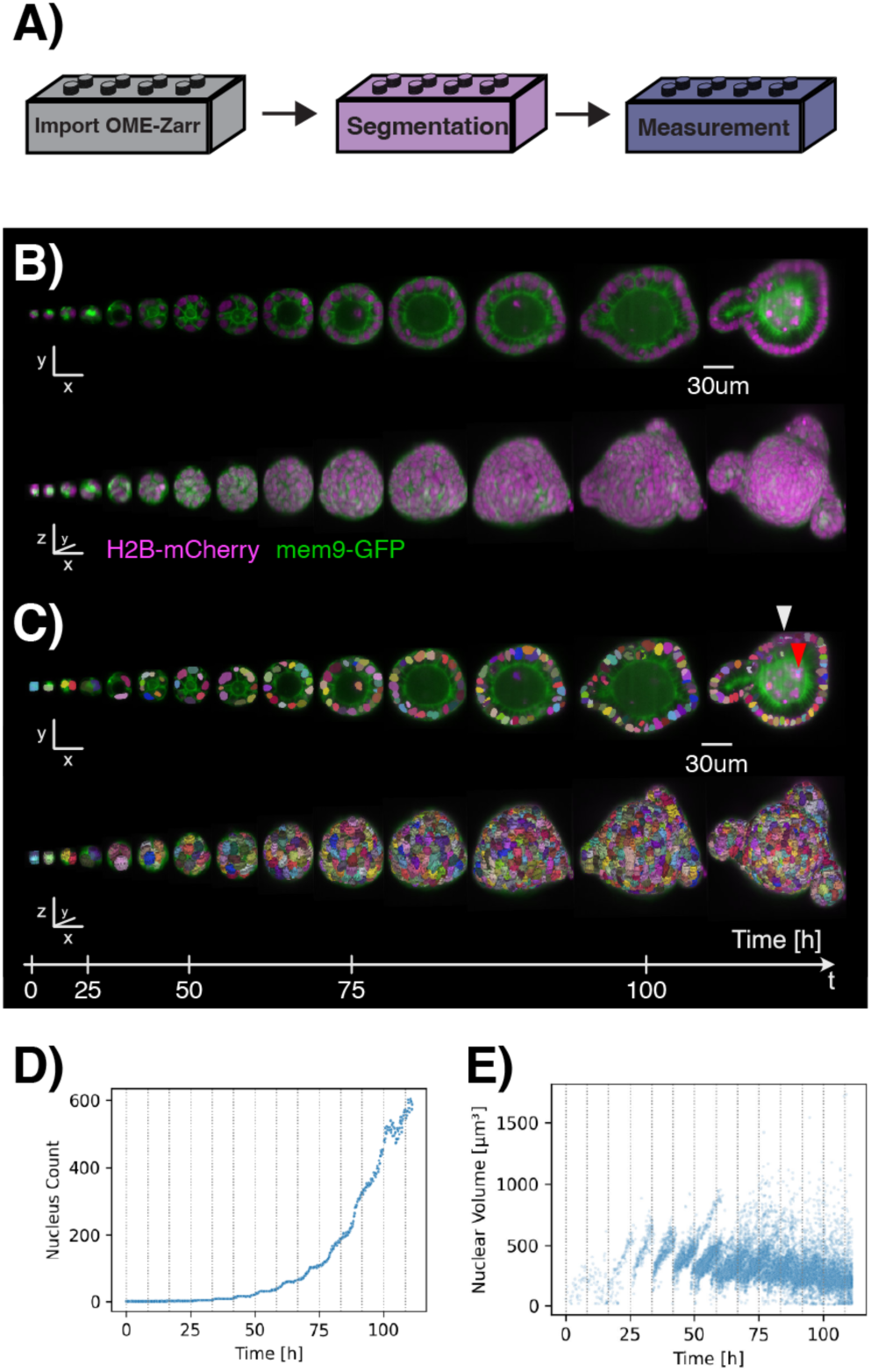
The Fractal task specification and the Fractal platform allow processing large 3D and time-resolved image data. **A** Overview of a simple Fractal workflow used for time series processing. **B** 2D (top) and 3D (bottom) rendering of a mouse intestinal organoid imaged over 110 hours of development, labelled with H2B-mCherry (magenta) and mem9-GFP (green). **C** 2D (top) and 3D (bottom) rendering of the segmentation results obtained with the cellpose SAM task. The white arrow indicates a mis-segmented nucleus in a late time point, while the red arrow indicates a false positive nucleus detection in the lumen. **D** Quantification of nuclei count over time. **E** Nuclear volume over time. Later time points were subsampled to 32 nuclei per time point to avoid overplotting.

### Clinical image analysis workflows

In our final example, we illustrate how the Fractal platform and OME-Zarr-based analysis workflows can be used to bring modular, interoperable image analysis workflows to clinical applications, focusing on an *ex vivo* study of patient-derived cells using microscopy-based readouts. High-throughput drug screening allows the identification of individualized susceptibilities of cancer cells to small-molecule perturbations and makes it possible to match the treatment of patients to the phenotypic characteristics of their individual cancer, an approach referred to as functional precision oncology^34^. Through this, microscopy image analysis can support the rational design of targeted interventions^35–37^.

At the University Children’s Hospital Zurich, we have established an image-based drug response profiling (DRP) platform for patients with acute leukemia, integrated in a European network of study centers and clinical trials^38,39^. In the DRP assay, bone marrow aspirates are co-cultured with mesenchymal stroma cells and exposed to a panel of about 100 drugs for 3 days before reading out the leukemia cell viability by automated microscopy. The computational analysis is separated into upstream image analysis using the Fractal platform and downstream modeling of dose responses, followed by cohort-based drug ranking (**Figure 6**). Phenotypic profiling unveils considerable heterogeneity in drug responses within and across leukemia subtypes, underscoring the utility of functional assays for therapeutic decision support. The open-source ecosystem of Fractal enables federated deployments that are compatible with in-house diagnostics developed at the hospital, thus effectively bringing the algorithms to the data, for the benefit of patients. Additionally, the platform ensures the reproducibility and traceability of the analysis pipelines, which is a prerequisite when translating drug screening results to patient care. The raw microscope output data is available on Zenodo^40^ and the processed OME-Zarr containers will be publicly available in the Bioimage Archive (*data submission in progress, link to be added in the final version of the manuscript*).

**Figure 6.**
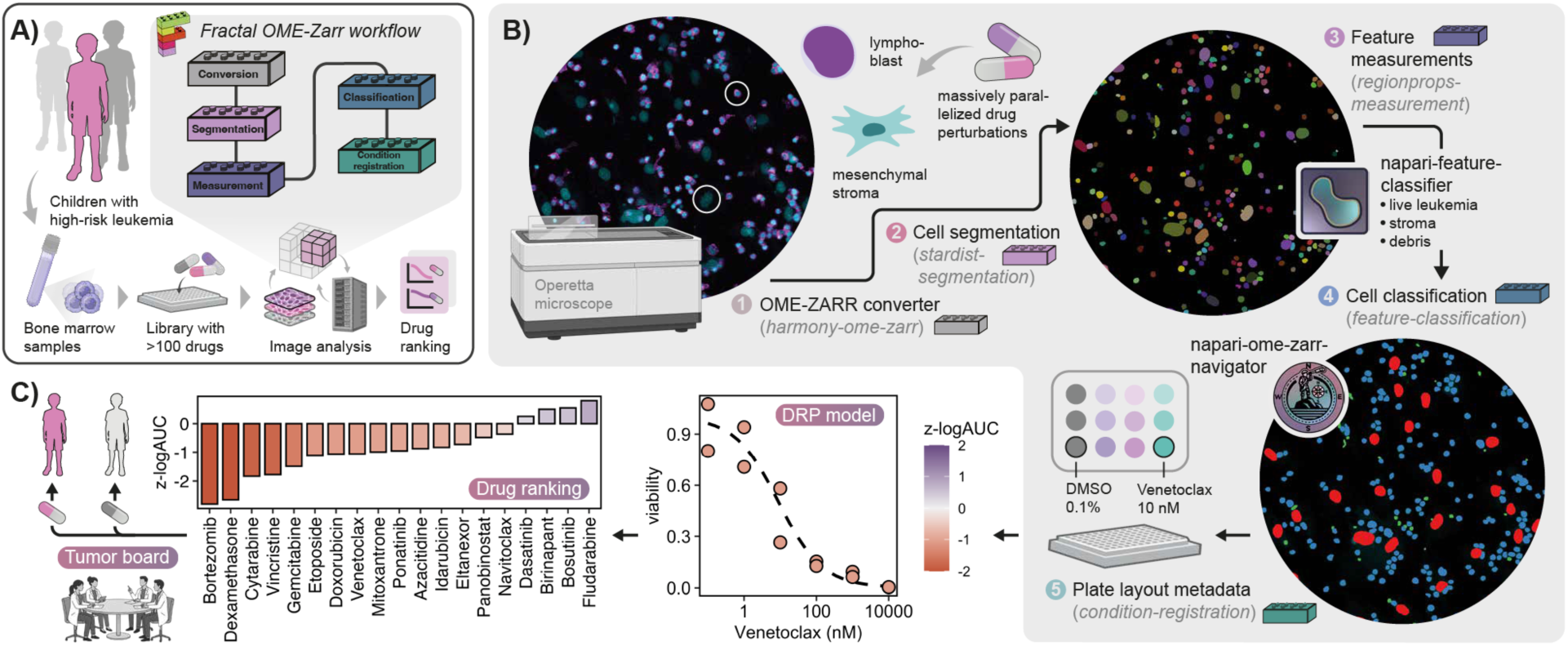
The Fractal task specification and the Fractal platform allow processing high-throughput screens for drug response profiling in hematological malignancies. **A** Cancer cells from leukemia patient derived xenograft were perturbed with a drug library and analyzed using a custom Fractal workflow. **B** We (1) converted images to OME-Zarr, (2) segmented individual nuclei with StarDist, (3-4) classified them by viability and cell type, and (5) annotated perturbation conditions on the plate. **C** We then counted viable cells as a function of drug concentration and normalized by a vehicle control (DMSO). Finally, we computed dose-response metrics for each drug and integrated across the patient cohort. Drug screening results were evaluated in the context of a tumor board. Figure prepared with icons from Biorender.

To demonstrate the reproducibility of Fractal workflows for drug profiling, we ran the workflow described above on 3 individual Fractal deployments and compared the obtained results. Each workflow consistently segmented and quantified 1,595,788 cells across the 240 wells and detected the exact same number of mesenchymal stem cells, leukemia cells and dead cells in each well, thus producing identical dose-response relationships. When comparing single-cell measurements for all cells across the different runs of the workflow and rounding numerical precision to 6 digits, more than 99.99% of cells obtained exactly matching measurements between all replicates. Mismatches in feature measurements between replicate runs only concerned 21-70 cells (0.001–0.004% of all objects) and were likely caused by minor segmentation boundary differences due to machine precision effects influencing the post-processing steps of StarDist (particularly non-maximum suppression and mask rasterization). This level of reproducibility across different infrastructure could be achieved by simply sharing the Fractal workflow files that fully specify the order of tasks to be run, their exact version number, and all their parameters.

## Discussion

Microscopy technologies continue to develop rapidly, offering systems with better resolution, faster acquisition, and richer multidimensional data, while concurrent breakthroughs in computer vision and AI are making it technically possible to extract quantitative information from these complex datasets. Together, these developments offer an unprecedented opportunity for biological discovery. However, the on-the-ground reality remains that most processing methods are developed for small images or narrow use cases, and that reusing them in a different context typically requires advanced programming skills. Working with large images further adds complications by introducing scientific computing challenges and effectively turning what would otherwise be straightforward analysis into software engineering endeavors. While these are difficult problems, the advent of standardized, scalable file formats like OME-Zarr offers a path forward. With a common data structure in place, we can begin to envision a future in which analysis methods no longer need to be reimplemented for each new context, and in which researchers acquiring data are empowered to design their own analyses by composing existing, validated processing modules. The core ingredient for realizing this vision is interoperability.

In this work, we proposed a concrete model towards that goal: a general definition for interoperable processing units built on OME-Zarr (the Fractal task specification) together with a demonstration of how flexible, scalable, and accessible computational frameworks can be constructed around it (the Fractal platform), with increased scalability as compared to interactive platforms such as Fiji^41^ and napari^6^. Through the diverse applications we presented, we illustrate the value of interoperable processing modules: they enable the design of tailored workflows addressing distinct biological questions, all from individual components that can operate both within and beyond the Fractal framework. This provides a first demonstration of how the OME-Zarr specification can be leveraged to harness HPC resources and deliver scalable yet user-friendly platforms that bring the community closer to FAIR bioimage analysis.

Compared to existing workflow-based image analysis software such as CellProfiler^42^ and KNIME^43^, the key strength of the Fractal platform lies in its native integration of the Fractal task specification, and therefore of the newly-developed OME-Zarr file format. This design choice greatly simplifies the handling of complex analysis problems, such as large volumetric data, big collections of fields of view to be processed in context, complex time-resolved datasets, or any combinations thereof. As the Fractal task specification is general and independent of the Fractal platform, the collection of available tasks can be augmented by any community contribution that follow the specification. While the Fractal platform demonstrates an integrated vision of OME-Zarr-centric image processing from task specification to user interface and viewer integration, the interoperability of the task specification also ensures that complex processing tasks developed for this framework can be reused with minimal adaptation in more general-purpose workflow orchestration systems that are already established.

We expect future developments in the OME-Zarr specification to significantly contribute to making the Fractal task specification and platform even more flexible. While ways to store collections of OME-Zarr with related metadata have already been defined for high-content screening data, further structured collection metadata will be essential to allow handling other kinds of images. As community work towards a collection specification for OME-Zarr has already started (https://ngff.openmicroscopy.org/rfc/8/index.html), we can expect a future version of the Fractal task specification and platform to adopt it, thereby facilitating the creation of more complex processing graphs, enabling more advanced parallelisation approaches, and simplifying interactions with read-only and remote input data. These developments will also facilitate the integration of processing modules following the Fractal task specification into workflow systems such as Galaxy, which come with stricter data staging requirements.

Although Fractal is currently primarily amenable to light microscopy, both the task specification and the platform are, thanks to their reliance on OME-Zarr, general enough to accommodate other imaging modalities, including electron microscopy and spatial transcriptomics, among many others. With substantial ongoing developments in analysis methods for these modalities, we anticipate the ecosystem of task packages to expand accordingly. Importantly, the open and community-driven nature of the task packages means that this expansion can occur organically, with or without our direct involvement, thus amplifying the potential of the platform for capabilities we do not yet foresee.

The past two decades have seen a remarkable growth of the bioimage analysis community, which has established itself as a vital bridge between computer science and biology. We are now at a point where the most pressing gap is no longer the availability of microscopy-specific computational methods, but their accessibility and reusability. Just as OME-Zarr is driving the standardization and unification of microscopy data management, we believe it can serve as the foundation for standardizing analysis methods and pipelines, ultimately empowering all life scientists to design and execute cutting-edge analyses themselves. Significant challenges remain, but the momentum of community-driven development, to which our work contributes, suggests that truly accessible, reproducible, and scalable microscopy image analysis has now become an achievable goal rather than a distant aspiration.

## Materials & methods

### The Fractal task specification

Working with structured image data in the OME-Zarr format increases the flexibility of processing for complex and large image structures. This is especially relevant when data are multi-dimensional and when multiple images must be processed in context, for instance, when hundreds of fields of view in a high-content screening plate must be tiled or stitched together. The chunked nature of the OME-Zarr format, combined with metadata on regions of interest that can span across original field of view boundaries, enables processing of arbitrary subsets of larger-than-memory datasets. While tools such as CellProfiler offer powerful analysis approaches when workflows can be run per field of view and cells at the edge of individual images can be excluded^42^, images of larger 3D structures, such as organoids or embryoids, often require processing across fields of view and a more flexible approach for handling images that are too large to be loaded in memory.

The Fractal table specification sets serialisation options, required metadata, and requirements for table contents. Fractal tables are added to an OME-Zarr container in an additional subgroup called “tables” that is similar to the “labels” subgroup. This association defines which image and labels a given table is associated with. The tables subgroup must contain Zarr attributes with a “tables” key and a list of table names as values. A corresponding Zarr group or folder must exist for each table name. Within an individual table folder, table contents are serialised as AnnData in the Zarr format^24^, Apache Parquet, CSV, or JSON, an implementation inspired by the way SpatialData uses AnnData tables^44^. Each table group must contain metadata specifying the *backend* used, as well as an *index_key* and *index_type*. Fractal tables are always typed with the *generic_table* type as a fallback for tables that do not follow specific restrictions on their content. Further technical details on table backends and on Fractal table types can be found at https://biovisioncenter.github.io/ngio/stable/table_specs/backend/ and https://biovisioncenter.github.io/ngio/stable/table_specs/overview/, respectively.

Feature tables contain measurements related to specific objects in label images, and have both metadata relating them to a given label image and an index column allowing to match the rows in the table with the corresponding objects in the label image. ROI tables contain named bounding box coordinates in physical units (and thus independent of the image resolution) to facilitate the loading of specific regions in a larger image. Masking ROI tables use segmentation labels to mask the underlying image and load only a specific subset of a rectangular bounding box. Condition tables allow associating tabular experimental metadata with OME-Zarr images, for instance, different treatment conditions or quality control readouts. Taken together, the table-ready OME-Zarr container thus represents a biologically meaningful unit of image data, derived segmentation, and quantifications.

The Fractal task specification supports five different types of tasks depending on the specificities of the OME-Zarr input and on parallelisation requirements:

1. <u>Parallel processing tasks</u> take a single OME-Zarr container as input (via a *zarr_url* parameter) and store their result either in the same OME-Zarr container (e.g., as new label subgroups or new tables) or in a new OME-Zarr container (e.g., a derived image). They can be run in an embarrassingly parallel fashion across a large list of OME-Zarr containers.
2. <u>Non-parallel tasks</u> receive a list of OME-Zarr containers as input (*zarr_urls* parameter) together with the directory for the Zarrs (*zarr_dir* parameter) and run a single process across all of them. These processing units store their outputs back into the OME-Zarr containers they received as input or in other files (e.g., quality control reports generated from many OME-Zarr containers may be stored at the plate level instead of in individual OME-Zarr containers). Non-parallel tasks are used when aggregating information across many OME-Zarr containers.
3. <u>Compound tasks</u> are used when dealing with more complex parallelisation requirements and consist of a non-parallel initialisation phase followed by a parallel compute phase. The initialisation phase processes a list of OME-Zarr inputs (*zarr_urls* parameter) together with the directory for the Zarrs (*zarr_dir* parameter) and generates a JSON configuration file for each parallel operation to be run. In that way, defined groups of OME-Zarr containers can be processed together, while separate parallel operations can also be run on the same OME-Zarr container.
4. <u>Converter tasks</u> are an exception to the rule that all tasks take OME-Zarr containers as inputs. As they generate OME-Zarr containers from microscopy images in other file formats (e.g., proprietary file formats or OME-tiff), they only take as input the directory where the resulting OME-Zarr containers should be written to (the *zarr_dir* parameter) as a required parameter. There are 2 types of converter tasks:

a. <u>non-parallel converters</u> run a single process to convert the input data to OME-Zarr;
b. <u>compound converters</u> use an initialisation phase akin to compound tasks to determine how many jobs should be started and carry initialisation arguments over to each compute task via JSON files.

Besides required input parameters such as *zarr_urls(s)* and *zarr_dir*, tasks can take an arbitrary number of additional parameters, which can either be mandatory or optional. Parameters are defined in the task manifest, which describe their specific requirements either as simple types (e.g., number, string, boolean, lists, tuples) or as complex, potentially nested models. All parameters are passed to a running task as a single JSON file through its CLI interface.

Tasks can also generate metadata that either specify newly created OME-Zarr containers and their associated metadata, or new metadata associated with existing OME-Zarr containers. These metadata are distinct from the OME-Zarr metadata that are directly written into the Zarr containers. To allow tracking newly generated OME-Zarr containers, the metadata JSON file outputted by a module implemented to follow the Fractal task specification must contain an *image_list_updates* list, which enumerates individual JSON objects with at minimum a *zarr_url* for every new OME-Zarr created. Additional metadata such as *origin* (e.g., the original source of a derived image), as well as type and attribute metadata, can also be provided by tasks in their image list updates. Further information on the Fractal task specification as well as illustrative examples can be found on the Fractal task specification documentation page at https://fractal-analytics-platform.github.io/tasks_spec/.

### Interoperability of Fractal tasks

The Fractal task specification defines a standardised command-line interface that all tasks must implement. Tasks accept parameters as a JSON file (--args-json) and output metadata path (--out-json) arguments. This interface is the key to interoperability, as any framework capable of running a shell command can orchestrate a Fractal task without modification to the task code. We demonstrate this with the *fractal-cellpose-sam-task* v0.1.9 applied to a publicly available test dataset^26^ and invoked in four ways: via the Python API (direct function call), via a bash script using the command-line interface entry point, via a Nextflow^20^ workflow, and via a Snakemake^21^ workflow.

Adapting the task for Nextflow and Snakemake required the implementation of minimal framework-specific glue code. For Nextflow, the JSON manifest (`_FRACTAL_MANIFEST_.json`), which ships with every Fractal task, contains a full JSON schema for all parameters (including types, descriptions, and defaults) that can be automatically converted to a Nextflow-compatible parameter template and to an *nf-schema-format schema* for *nf-core* compatibility. The Fractal parallel task model (i.e., one invocation per OME-Zarr image) maps naturally to a Nextflow channel, where each *zarr_url* becomes an independent process execution. For Snakemake, the same parallelism is expressed through an integer index wildcard over the list of *zarr_url*, with task parameters supplied via a YAML config file. In both cases, only the required parameters need to be specified explicitly as all others fall back to the manifest defaults of the task.

All four examples were managed with pixi and are reproducible with a single pixi run command per example. The full code, configuration files, and setup instructions are available online at https://github.com/fractal-analytics-platform/fractal-task-interoperability.

### The Fractal platform

Workflow management is required to scale up image processing for larger datasets and to reproducibly execute complex series of tasks. In order to run workflows on terabytes-scale data, both computational resources and the data to be processed must be co-located as typically offered in HPC environments. HPC environments, however, come with added complexity: they typically require users to be comfortable with command line usage and to understand schedulers such as Slurm^45^. As a result, the accessibility barrier is often too high for researchers with limited computational expertise, and significant training efforts are required to get onboarded to HPC systems. To address this, we propose an abstraction layer on the usage of HPC clusters to ensure that life scientists can perform FAIR bioimage analysis with minimal technical overheads.

### Main components

The Fractal platform is not a centralized service: it is deployed on specific computational environments and is then available to the users having access to them. The backend server of the Fractal platform allows users with access to the environment to orchestrate the submission of workflows to available compute resources, keep track of their detailed processing history, and install complex computational environments for the processing tasks. To ensure scalability, it is built as a FastAPI app (https://fastapi.tiangolo.com/) that provides the server interface. The FastAPI app talks to a PostgreSQL database (https://www.postgresql.org/) with defined table schemas for users, workflows, datasets, processing histories, among other functionalities. This allows the server to expose a Representational State Transfer (REST) API and to scale easily with many users managing their projects, configuring their workflows, submitting and monitoring jobs, and installing task packages. While a well-optimised computer in a given lab may be able to perform database operations such as managing projects and workflows, the actual analysis of terabyte-scale image datasets requires the computational resources offered by institutional HPC clusters. To scale the analysis on available compute resources, the Fractal platform therefore typically connects to Slurm clusters (https://slurm.schedmd.com/), either by running the Fractal backend server on a (secondary) Slurm login node or by connecting to Slurm via ssh. All scientific data as well as computational task environments, are stored in the HPC cluster space. The backend Fractal server can trigger the automated installation of task packages from PyPI (https://pypi.org/), Pixi configurations (https://pixi.prefix.dev/), or Python wheel files. Alternatively, system administrators can manually create Linux command-line interface executables and register them with the backend Fractal server.

The actual processing workflows are submitted by the Fractal platform to the available computational resources using one of the modular Fractal runners, which offer an interface between the Fractal backend server and the HPC cluster architecture. We provide custom Fractal runners for Slurm as well as for local execution, the latter being primarily targeted at testing and running small workflows. The runners receive workflows to be executed and manage their orchestration, reporting activity and progress back to the backend Fractal server. In that manner, a single backend server can moderate dozens of parallel workflows running on an HPC cluster and scale the resource usage automatically.

To allow users with limited computational expertise to interact with the Fractal backend server and to benefit from the abstractions it provides, we provide easy access through the Fractal web interface (https://github.com/fractal-analytics-platform/fractal-web), a web application based on the Svelte UI framework (https://svelte.dev/) and served through a Node.js server (https://nodejs.org/) which provides an accessible web interface for users to interact with the backend server REST API. The Fractal web application and backend server integrate with Open Authorization (OAuth) identity providers to allow institutional single sign-on. The web interface offers an overview of projects, tasks, and jobs, as well as an integration with data visualisation tools. Users manage their datasets and workflows in the Fractal platform through so-called *Projects*, while *Datasets* keep track of any OME-Zarr created by a workflow and its metadata and are used by workflows to decide which OME-Zarrs to process. *Workflows* are defined as a series of processing operations following the Fractal task specification, and the Fractal web interface exposes all advanced workflow submission options. Users can run whole or partial workflows, and can run a given workflow on a full dataset or on sub-selections thereof. The Fractal web interface integrates with the backend Fractal server history models of workflow runs to enable the display of simple summary overviews of the processing status of all the images, while also exposing advanced access to detailed execution logs reporting on task execution and selected parameters. On top of the project and workflow management functionality, the Fractal web interface also contains a *Task page* and a *Jobs page*. On the task page, users can explore all the task packages that have been installed on their Fractal deployment and trigger the installation of additional task packages. For task packages built following the Fractal task specification, the complex Python environment installation is thereby simplified into a no-code experience. On the jobs page, users can get an overview of all currently running or previously run workflows, and have access to the same detailed logs as on the workflow page.

For advanced users and to enable automation, we also provide command-line-based access to most of the functionality above. The Fractal client (https://github.com/fractal-analytics-platform/fractal-client) exposes the REST-API of the Fractal backend server directly on the command line.

### Data service

The Fractal data service offers HTTP streaming of on-disk OME-Zarr datasets, enabling web-based access to data stored on HPC storage systems. Read access to on-disk data is implemented either by direct mounting of the HPC storage, or via the SSH/SFTP protocol (notably as implemented in the rclone-mount tool, https://rclone.org/commands/rclone_mount). User authentication for the data service relies on the backend server to allow integration with institutional single sign-on. A granular access-control layer, implemented across the backend server and the data service, guarantees that only authorized users can stream OME-Zarr data. This includes the user who created a given dataset, and any user who was granted access to it by the original creator.

### Napari plug-ins

Although existing napari plugins like *napari-ome-zarr* (https://github.com/ome/napari-ome-zarr) already allow the dynamic streaming of OME-Zarr image data, we developed a new suite of plugins to offer additional functionalities. First, the *napari-ome-zarr-navigator* plugin (https://github.com/fractal-napari-plugins-collection/napari-ome-zarr-navigator) facilitates the interactive visualization of selected regions of interests in images, labels, and tabular data in OME-Zarr containers that are available either locally (e.g., stored directly on a local hard drive or accessible on a mounted share) or remotely through the authenticated Fractal data service. It offers an interface to explore the image data and integrates with potential condition tables stored in the OME-Zarr container. The plugin also implements a ROI functionality that loads specific regions of images or labels at a given resolution into memory as napari layers, based on the ROI tables available in the OME-Zarr. This enables easier interactivity with other existing napari plugins that do not support multi-resolution layers and better 3D visualisations for parts of large 3D volumes without the need for lazy loading. The ROI loader can also read feature tables stored in the OME-Zarrs and attach them to napari label layers. Loading image layers, label layers, and associated measurements into napari makes it possible to interactively visualize processing results and simplifies further local processing in napari. Second, the *napari-feature-visualization* plugin (https://github.com/fractal-napari-plugins-collection/napari-feature-visualization) colors label objects according to pre-computed measurements. Third, the *napari-feature-classifier* plugin (https://github.com/fractal-napari-plugins-collection/napari-feature-classifier) facilitates the iterative training of random forest object classification models.

### Cardiac differentiation multiplexed image analysis workflow

We developed a *Convert Cellvoyager Multiplexing to OME-Zarr* conversion task, released together with other tools in the *fractal-tasks-core* task package (https://github.com/fractal-analytics-platform/fractal-tasks-core) to convert input images from TIFF to OME-Zarr. Each well of the multiwell plate was saved as a single larger tiled OME-Zarr image per cycle, combining all fields of view into a single array based on their stage metadata. As a result, plate visualisation is simplified, and the number of image pyramids and individual chunk files is heavily reduced. We also saved ROI tables with the original fields of view to make processing per field of view in downstream tasks easier. For the pre-processing steps, we first developed an *Illumination Correction* task (packaged in *fractal-tasks-core*) to correct the flatfield illumination bias inherent to the image acquisition process. It first subtracts the average background intensity value and then adjusts image intensities relying on a pre-calculated flatfield image constructed from a smoothed averaging of the per-channel XY planes. For the multiplexing registration, we used the *Elastix registration* tasks developed as part of the *abbott* Fractal tasks package (https://github.com/pelkmanslab/abbott). We first calculated the required B-spline transformation for each field of view between each cycle and the acquisition 0 reference cycle using the *Compute Registration (elastix)* task. We then applied the registration to each cycle with the *Apply Registration (elastix)* task, generating a new registered OME-Zarr image for each well and multiplexing acquisition. We carried out qualitative quality control checks for the registration by visually inspecting the alignments in projected plate overviews in order to identify fields of view with major registration issues. The main causes of registration issues were either parts of the sample detaching during multiplexing acquisitions or experimental handling. For fields of view that remained physically intact, registration was observed to be satisfactory both in the XY plane as well as in orthogonal (XZ and YZ) planes. This registration operation was found to correct both for smaller sample deformation in 3D over the experiment, as well as for minor drifts of the microscope, leading to a small unavailable area at the edge of each field of view (which we set to 0 and ignored during downstream analysis). To avoid data duplication, only the OME-Zarr images that had been both illumination corrected and registered were kept for further analysis.

Once illumination-corrected and aligned, we segmented individual 3D nuclei using the pretrained cyto2 model of cellpose 2^46^ wrapped in the *Cellpose Segmentation* task we developed as part of *fractal-tasks-core*. The limiting factors for the 3D nuclear segmentation quality were the axial resolution (1μm) as well as the presence of imaging and staining artefacts in dense 3D regions. While membrane stains were available in this dataset, we did not pursue cell segmentation as the expression level of the various membrane markers exhibited too strong variations across the differentiation time course. We then extracted 3D measurements for the reference cycle using the *scMultiplex Feature Measurement* task developed as part of the *scMultiplex* package (https://github.com/fmi-basel/gliberal-scMultipleX), which implements 3D morphological quantification (e.g., object surface areas) and intensity measurements for anisotropic images.

As the imaged cells form a pseudo-2D structure and only occasionally assemble into 2-3 cell layers in height, the dataset could also be analyzed in 2D. We therefore also calculated the maximum intensity projection of the image volume using the *Project Image (HCS Plate)* task we developed as part of *fractal-tasks-core*, and segmented nuclei with the cyto2 model wrapped in the same *Cellpose Segmentation* task mentioned previously. We finally filtered the segmented nuclei with the *Filter Label by Size* task developed as part of the *APx Fractal task collection* package (https://github.com/pelkmanslab/APx_fractal_task_collection), extracted features relying on the *Measure Features* task from the *APx Fractal task collection*, and aggregated these measurements across all multiplexing cycles using the *Aggregate Feature Tables* task from the *APx Fractal task collection*. These features could then be used to carry out object classification for quality control and cell type classification using the *napari-feature-classifier* (see **Napari plug-ins** subsection above) and the *Feature Classification* task from the *operetta-compose* package (https://github.com/leukemia-kispi/operetta-compose). We downloaded the 2D feature measurements across all wells, and filtered approximately a third of the originally segmented 1.2 million objects for quality control reasons. These included registration issues, staining artefacts, segmentation errors (as detected by classifiers and using the Fractal feature explorer), and the position of objects close to an edge of a measurement region. The filtered measurements were combined with classification results and metadata about the day of cardiac differentiation stored in the condition table of the dataset.

The full Fractal workflow is available in the paper Github repository (https://github.com/fractal-analytics-platform/fractal-paper-2026) under `fractal_paper_2026/workflows/cardiac_diff_workflow.json`, along with the code used for quality control, UMAP projection, and plotting.

### Zebrafish embryos 3D-4i image analysis workflow

We first converted the raw microscopy data into OME-Zarr using the *Convert Cellvoyager Multiplexing to OME-Zarr* task from *fractal-tasks-core*. Following this, we corrected uneven microscope illumination by estimating and applying flatfield illumination correction models per wavelength irreversibly to the images using *Calculate BaSiCPy Illumination Models* (to estimate the bias on separate reference images) and *Apply BaSiCPy Illumination Models*, two tasks packaged in the *APx Fractal task collection*. Next, to correct for pixel-shift in multi-channel acquisitions, we used *abbott*’s *Compute/Apply Channel Registration (elastix)* task and estimated a similarity transform and registration to a reference channel. We subsequently corrected sample shift and shrinkage/expansion across multiplexing cycles with the *Compute/Apply Registration (elastix)* task (also packaged in *abbott*), which estimated and applied three sequential transforms (rigid, affine, and B-spline), each step introducing more degrees of freedom to achieve a high-quality alignment at the single-cell level. We assessed the quality of the registration with the *Calculate Cycle Registration Quality* task of the *abbott-feature* package (https://github.com/pelkmanslab/abbott-features), and excluded structures with a low registration quality from downstream analysis. To segment multicellular structures, we trained a pixel classifier in the GUI of the open-source software Ilastik^7^, and used the trained model on the whole data through the *Ilastik Pixel Classification Segmentation* task of the *fractal-ilastik-tasks* package (https://github.com/fractal-analytics-platform/fractal-ilastik-tasks). For nuclei segmentation, we used the *Cellpose Segmentation* task from *fractal-tasks-core*, and for individual cell segmentation we relied on *abbott*’s *Seeded Watershed Segmentation* task using the previously generated nuclei labels used as seeds. Feature extraction was performed using the *abbott-features* package: intensity decay models were first estimated to correct for time-dependent signal loss (using the *Get Cellvoyager Time Decay* task) and z-intensity decay from spherical aberration and scattering of light in the tissue (using the *Get Z Decay Models* task). Finally, systematic multiscale feature extraction was performed by extracting the (corrected) features using the *Measure Features* task. We here compared the cell type classification annotations with the Leiden clusters using the *napari-feature-classifier* (see **Napari plug-ins** subsection above) outputs from the original study^30^, although the resulting random forest classifiers could also be used directly in Fractal using the *Feature classification* task from the *operetta-compose* package.

The full Fractal workflow associated to these results is available in the paper Github repository (https://github.com/fractal-analytics-platform/fractal-paper-2026) under `fractal_paper_2026/workflows/workflow-temp_3d_multiplexed_abbott.json`, along with the code to generate the figure.

### Time-lapse volumetric image analysis workflow

We first tuned the resolution and pre-process blurring parameters of the *Cellpose SAM Segmentation* task in the Fractal web interface on a subset of the time-lapse frames by employing the ROI table interface to specify a subset of data to be processed. The final workflow was then applied on images downsampled by a factor of two in X and Y due to anisotropic sampling at the highest resolution level (Z=2.0µm, X and Y=0.406µm), without pre-process blurring. A significant speedup (∼10x) in processing time was achieved by specifying a ROI table containing tight bounding boxes around the organoid in each frame. The same tables could be re-used to generate the visualizations in **Figure 5B-C**. The ROI tables were computed in a separate script (`fractal_paper_2026/light_sheet_analysis/compute_bounding_boxes.py`) outside of Fractal. To quantify nuclear counts and nuclear volume, we used the *Region Props Features* task from the *fractal-tasks-template* package (https://github.com/fractal-analytics-platform/fractal-tasks-template).

The full Fractal workflow associated to these results is available in the paper Github repository (https://github.com/fractal-analytics-platform/fractal-paper-2026) under `fractal_paper_2026/workflows/light_sheet_workflow.json`, along with the code to generate the figure.

### Drug profiling image analysis workflow

We developed a dedicated *operetta-compose* task package to build a workflow structured as follows. First, raw tiff images of live cells recorded on a high-content microscope (Operetta, Revvity) with channels for viability (CyQuant) and optionally cell surface markers (e.g., CD19, CD7) were converted into OME-Zarr relying on the tailored *Harmony to OME-Zarr* conversion task of the *operetta-compose* package developed for this workflow (https://github.com/leukemia-kispi/operetta-compose). Cell nuclei were then segmented with StarDist relying on the *Stardist segmentation* task, and intensity and label features were measured using the *Regionprops measurement* task wrapping scikit-image’s regionprops method^47^. We then trained a random forest classifier in napari using the *napari-feature-classifier* plugin (see **Napari plug-ins** subsection above) to distinguish mesenchymal stromal cells from live leukemia cells and applied it to the whole dataset using the *Feature classification* task. Plate metadata, including well positions of drugs and their concentrations, was registered as a Fractal condition table using the *Condition registration* task. Leveraging the *napari-ome-zarr-navigator* plugin (see **Napari plug-ins** subsection above), we could interactively explore conditions, images, and ROIs in the OME-Zarr dataset. Finally, we fitted log-logistic dose response models and computed metrics such as the logarithmic half-maximal effective dose (logEC50) and the area-under-the curve (logAUC). These parameters were integrated across the patient cohort to rank drugs by sensitivity. In the tumor board, individualized therapeutic interventions were prioritized based on clinical, molecular, and functional data points.

For the reproducibility analysis, we did not carry out the comparison of measurements based on matching labels, as the labels assigned by StarDist can vary. Instead, we rounded all numerical measurements to 6 digits of numerical precision and hashed the feature vector for each cell. Within each well, unique matches were assigned between matching hashes, while the feature vectors for non-matching hashes were saved for downstream evaluation to diagnose the source of variation.

The full Fractal workflow associated to these results (`fractal_paper_2026/workflows/drp_workflow.json`) as well as the source code for the dose-response calculation, figure generation, and the reproducibility analysis are available in the Github repository of the paper (https://github.com/fractal-analytics-platform/fractal-paper-2026).

## Acknowledgements

JL was supported by Chan Zuckerberg Initiative DAF, an advised fund of Silicon Valley Community Foundation, Grant #2021-240441 and Grant #2022-252401, by swissuniversities (BioFAIR, Projektvertrag PgB_25-28_674_A1_30). LC was supported by swissuniversities (BioFAIR, Projektvertrag PgB_25-28_674_A1_30). MH was supported by Allen Institute for Cell Science. RH was supported by the Promoter Foundation of the Department of Molecular Life Sciences at the University of Zurich. GQGM was supported by the Friedrich Miescher Institute for Biomedical Research (FMI). NAR was supported by the European Union’s Horizon Europe research and innovation programme under the Marie Skłodowska-Curie grant agreement No 101032127. LKC was supported by the Heidi-Ras foundation (PI: FDS). FDS was supported by Zurich Precision Oncology for Children (ZPOC), an UMZH technology initiative. FDS and JPB kindly acknowledge support from the University of Zurich, the Children’s Research Center (FZK) of the University Children’s Hospital Zurich and various foundations. LP was supported by the Swiss National Science Foundation (grants 320030-231576 and 320030-236346), the European Research Council (advanced grant CROSSINGSCALES-885579), and the University of Zurich. PL was supported by the Swiss National Science Foundation (grants 310030_208162 and TMCG-3_223263), the Allen Institute for Cell Science, and the Biohub.

The authors gratefully acknowledge members of the eXact lab for their contributions to the software engineering and technical implementation of the Fractal platform, in particular Marco Franzon, Yuri Chiucconi, and Sonia Zorba. Further gratitude extends to additional past and present contributors from eXact lab and the broader Fractal community who have supported the design, implementation, and testing of the platform. The authors also thank the community of developers of Fractal task packages and analysis components that extend the Fractal ecosystem, in particular Flurin Sturzenegger, Enrico Tagliavini, Marvin Albert, and other community contributors who continue to develop and share tasks following the Fractal task specification. Finally, the authors acknowledge their collaborators and institutional partners who provided scientific, technical, and infrastructure support. Particular thanks are addressed to the Facility for Advanced Imaging and Microscopy at FMI, especially Jan Eglinger, Tim-Oliver Buchholz, Sabine Reither, and Laurent Gelman; the Center for Microscopy and Image Analysis at the University of Zurich, especially Flurin Sturzenegger, Joana Martins, and Urs Ziegler; and the IT and infrastructure teams at FMI IT, UZH Science IT, UZH Central IT, and the UZH Department of Molecular Life Sciences IT Support for enabling and supporting Fractal deployments and high-performance computing infrastructure.

## Supplementary Figures

**Supplementary Figure 1.**
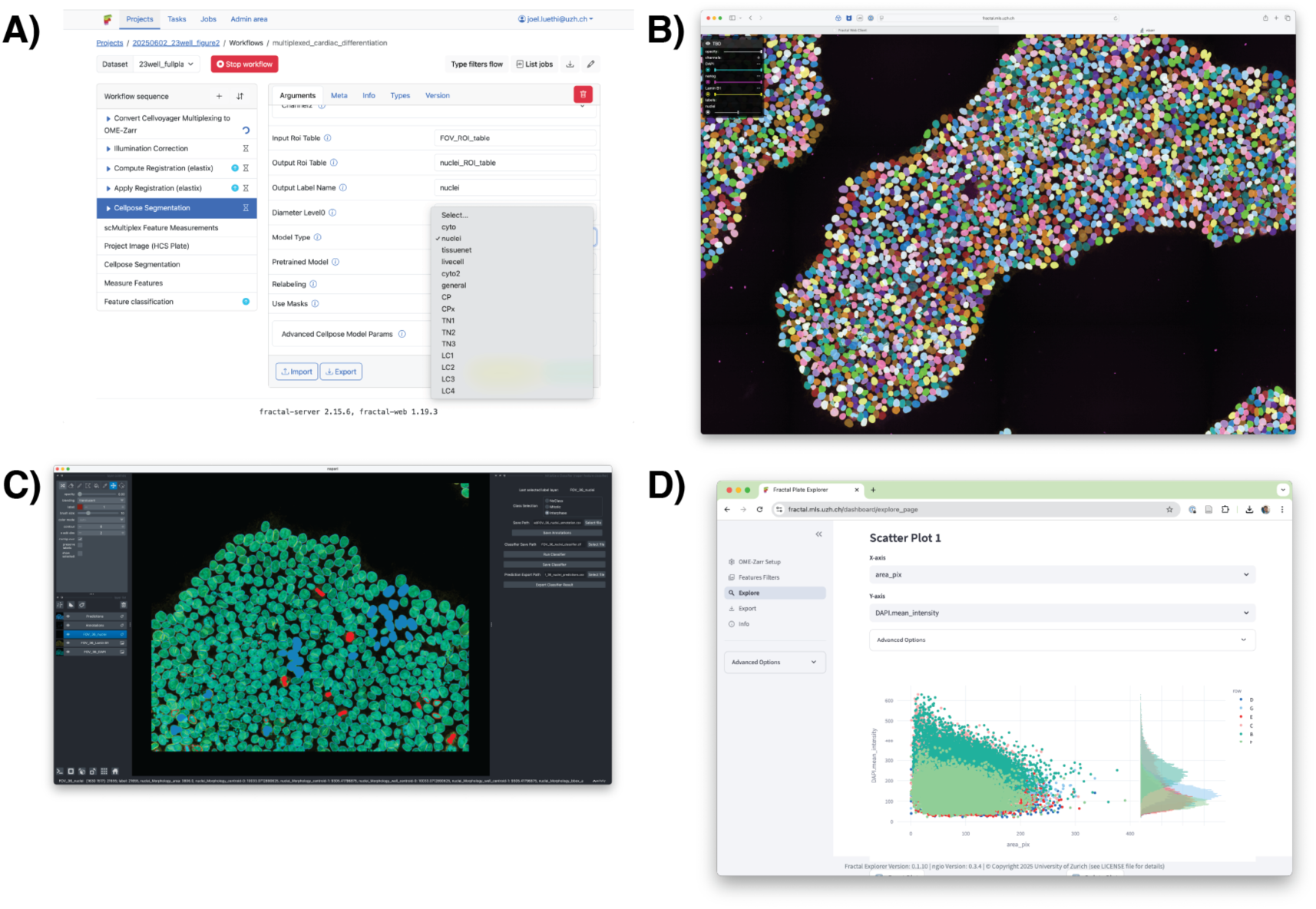
The Fractal platform facilitates the design and execution of complex image analysis workflows through a no-code web interface. **A** To demonstrate that the Fractal task specification can serve as a basis for accessible interoperable image processing relying on the OME-Zarr format, we have developed a web-based user interface and integrated it with existing OME-Zarr viewers for image visualisation. Workflow design, execution, and monitoring, as well as parameter tuning, can be carried out directly in the web interface, while the processing itself happens on a remote HPC. **B** To ensure that remote data remains accessible, we developed a data service that streams OME-Zarr data from HPC cluster storage over HTTP. This is integrated with the ViZarr web viewer^15^ to provide direct visualisation access in a web browser. **C** More advanced interaction with the image data, such as loading a partial subset of images or creating annotations to train classification models, is facilitated by a set of napari plugins we developed to stream OME-Zarr files and interact with them. **D** As the OME-Zarr containers also contain all the associated tabular measurement data, we also developed a web dashboard that allows for interactive access, filtering, plotting, and export of these measurements.

## References

1. Weigert, M. et al. Content-aware image restoration: pushing the limits of fluorescence microscopy. Nat Methods 15, 1090–1097 (2018).

2. Stringer, C., Wang, T., Michaelos, M. & Pachitariu, M. Cellpose: a generalist algorithm for cellular segmentation. Nat Methods 18, 100–106 (2021).

3. Umarov, R., Li, Y. & Arner, E. DeepCellState: An autoencoder-based framework for predicting cell type specific transcriptional states induced by drug treatment. PLoS Comput Biol 17, e1009465 (2021).

4. Palma, A., Theis, F. J. & Lotfollahi, M. Predicting cell morphological responses to perturbations using generative modeling. Nat Commun 16, 505 (2025).

5. Gómez-de-Mariscal, E. et al. DeepImageJ: A user-friendly environment to run deep learning models in ImageJ. Nat Methods 18, 1192–1195 (2021).

6. Sofroniew, N., et al. napari: a multi-dimensional image viewer for Python. Zenodo 10.5281/ZENODO.3555620 (2026).

7. Berg, S. et al. ilastik: interactive machine learning for (bio)image analysis. Nat Methods 16, 1226–1232 (2019).

8. Dalle Nogare, D., Hartley, M., Deschamps, J., Ellenberg, J. & Jug, F. Using AI in bioimage analysis to elevate the rate of scientific discovery as a community. Nat Methods 20, 973–975 (2023).

9. Ellenberg, J. et al. A call for public archives for biological image data. Nat Methods 15, 849–854 (2018).

10. Wilkinson, M. D. et al. The FAIR Guiding Principles for scientific data management and stewardship. Sci Data 3, 160018 (2016).

11. Goldberg, I. G. et al. The Open Microscopy Environment (OME) Data Model and XML file: open tools for informatics and quantitative analysis in biological imaging. Genome Biol 6, R47 (2005).

12. Moore, J. et al. OME-NGFF: a next-generation file format for expanding bioimaging data-access strategies. Nat Methods 18, 1496–1498 (2021).

13. Moore, J. et al. OME-Zarr: a cloud-optimized bioimaging file format with international community support. Histochem Cell Biol 160, 223–251 (2023).

14. Moore, J. et al. Enabling Peta-Scale Federated Repositories through Cloud-Native Formats: Lessons from a fast-paced challenge in the bioimaging community. https://doi.org/10.5281/ZENODO.16735915 (2025) doi:10.5281/ZENODO.16735915.

15. Manz, T. et al. Viv: multiscale visualization of high-resolution multiplexed bioimaging data on the web. Nat Methods 19, 515–516 (2022).

16. Pape, C. et al. MoBIE: a Fiji plugin for sharing and exploration of multi-modal cloud-hosted big image data. Nat Methods 20, 475–476 (2023).

17. Gros, O. et al. Microscopy Nodes: versatile 3D microscopy visualization with Blender. Preprint at 10.1101/2025.01.09.632153 (2025).

18. Barbiero, S., Soneson, C., Liberali, P. & Stadler, M. B. ez-zarr: A Python package for easy access and visualisation of OME-Zarr filesets. JOSS 10, 7882 (2025).

19. Özdemir, B., et al. BatchConvert: A command-line tool for parallelised conversion of image collections into the standard bioimage file formats OME-TIFF and OME-Zarr. Preprint at 10.3897/arphapreprints.e116669 (2023).

20. Di Tommaso, P. et al. Nextflow enables reproducible computational workflows. Nat Biotechnol 35, 316–319 (2017).

21. Mölder, F. et al. Sustainable data analysis with Snakemake. F1000Res 10, 33 (2025).

22. The Galaxy Community et al. The Galaxy platform for accessible, reproducible, and collaborative data analyses: 2024 update. Nucleic Acids Research 52, W83–W94 (2024).

23. Newman, D. J. Zarr Storage Specification. https://doi.org/10.5067/DOC/ESCO/ESDS-RFC-048V1 (2024) doi:10.5067/DOC/ESCO/ESDS-RFC-048V1.

24. Virshup, I., Rybakov, S., Theis, F. J., Angerer, P. & Wolf, F. A. anndata: Access and store annotated datamatrices. JOSS 9, 4371 (2024).

25. Pachitariu, M., Rariden, M. & Stringer, C. Cellpose-SAM: superhuman generalization for cellular segmentation. Preprint at 10.1101/2025.04.28.651001 (2025).

26. Lüthi, J. et al. 2024 OME-NGFF workflows hackathon. Preprint at 10.37044/osf.io/5uhwz_v1 (2025).

27. Gut, G., Herrmann, M. D. & Pelkmans, L. Multiplexed protein maps link subcellular organization to cellular states. Science 361, eaar7042 (2018).

28. Kramer, B., Castillo, J., Pelkmans, L. & Gut, G. Iterative Indirect Immunofluorescence Imaging (4i) on Adherent Cells and Tissue Sections. BIO-PROTOCOL 13, (2023).

29. Gerbin, K. A. et al. Cell states beyond transcriptomics: Integrating structural organization and gene expression in hiPSC-derived cardiomyocytes. Cell Systems 12, 670–687.e10 (2021).

30. Hess, M. et al. Multiplexed embryo profiling links cellular state to zygotic genome activation in single cells. Preprint at 10.1101/2025.10.27.684723 (2025).

31. McInnes, L., Healy, J., Saul, N. & Großberger, L. UMAP: Uniform Manifold Approximation and Projection. JOSS 3, 861 (2018).

32. Traag, V. A., Waltman, L. & Van Eck, N. J. From Louvain to Leiden: guaranteeing well-connected communities. Sci Rep 9, 5233 (2019).

33. De Medeiros, G. et al. Multiscale light-sheet organoid imaging framework. Nat Commun 13, 4864 (2022).

34. Letai, A., Bhola, P. & Welm, A. L. Functional precision oncology: Testing tumors with drugs to identify vulnerabilities and novel combinations. Cancer Cell 40, 26–35 (2022).

35. Lee, S. et al. High-throughput identification of repurposable neuroactive drugs with potent anti-glioblastoma activity. Nat Med 30, 3196–3208 (2024).

36. Kornauth, C. et al. Functional Precision Medicine Provides Clinical Benefit in Advanced Aggressive Hematologic Cancers and Identifies Exceptional Responders. Cancer Discovery 12, 372–387 (2022).

37. Malani, D. et al. Implementing a Functional Precision Medicine Tumor Board for Acute Myeloid Leukemia. Cancer Discovery 12, 388–401 (2022).

38. Frismantas, V. et al. Ex vivo drug response profiling detects recurrent sensitivity patterns in drug-resistant acute lymphoblastic leukemia. Blood 129, e26–e37 (2017).

39. Steffen, F. D. et al. Drug Response Profiling Informs Personalized Bridging to Cell Therapy for Patients with Relapsed/Refractory Acute Lymphoblastic Leukemia. Blood 142, 4350–4350 (2023).

40. Steffen, F. & Cantoni, L. Operetta Drug Profiling benchmark 10x multiwell plate. Zenodo 10.5281/ZENODO.18417467 (2026).

41. Schindelin, J., et al. Fiji: an open-source platform for biological-image analysis. Nat Methods 9, 676–682 (2012).

42. Stirling, D. R. et al. CellProfiler 4: improvements in speed, utility and usability. BMC Bioinformatics 22, 433 (2021).

43. Berthold, M. R. et al. KNIME: The Konstanz Information Miner. in Data Analysis, Machine Learning and Applications (eds Preisach, C., Burkhardt, H., Schmidt-Thieme, L. & Decker, R.) 319–326 (Springer Berlin Heidelberg, Berlin, Heidelberg, 2008). doi:10.1007/978-3-540-78246-9_38.

44. Marconato, L. et al. SpatialData: an open and universal data framework for spatial omics. Nat Methods 22, 58–62 (2025).

45. Yoo, A. B., Jette, M. A. & Grondona, M. SLURM: Simple Linux Utility for Resource Management. in Job Scheduling Strategies for Parallel Processing (eds Feitelson, D., Rudolph, L. & Schwiegelshohn, U.) vol. 2862 44–60 (Springer Berlin Heidelberg, Berlin, Heidelberg, 2003).

46. Pachitariu, M. & Stringer, C. Cellpose 2.0: how to train your own model. Nat Methods 19, 1634–1641 (2022).

47. Van Der Walt, S. et al. scikit-image: image processing in Python. PeerJ 2, e453 (2014).

